# Autophagy dependent HIF1α proteostasis is compromised in models of PEX1 deficiencies

**DOI:** 10.64898/2026.09.29.755113

**Authors:** Soham P. Chowdhury, Sambhav Jain, Milagros Esmerode, Henry Dong, Jonathan T. Vu, Connor J. Sheedy, Chris D. Richardson, Brooke M. Gardner

## Abstract

Peroxisomes, along with mitochondria, coordinate and compartmentalize oxidative metabolism in eukaryotic cells. Rare genetic disorders caused by mutations in PEX genes impair peroxisome function and cause Peroxisome Biogenesis Disorders, which are characterized by liver and neurological dysfunction, hearing and vision loss, and metabolic abnormalities. The majority of Peroxisome Biogenesis Disorders (PBDs) are caused by mutations in the gene encoding PEX1, which together with PEX6 forms a hetero-hexameric AAA-ATPase complex that drives the import of enzymes into the peroxisome lumen. One particular destabilizing mutation - PEX1^G843D^ - results in almost 30% of all cases. Here we show that deficiencies in PEX1 lead to increased levels of the oxygen-responsive transcription factor HIF1α in normoxia, as well as a HIF1α transcriptional signature. The increase in HIF1α protein was rescued by overexpression of PEX1^WT^, suggesting PEX1 deficiencies modulate HIF1α signaling. The increased levels of HIF1α were not explained by defects in the oxygen responsive degradation pathway of HIF1α that relies on proline hydroxylase domain enzyme-dependent hydroxylation and von Hippel Lindau tumor suppressor protein-dependent ubiquitination. Instead, we found that PEX1 deficiency alters HIF1α proteostasis by reducing degradation through a hydroxylation- and autophagy-dependent mechanism. Notably, enhancing autophagic capacity by ULK1 agonism was sufficient to reduce HIF1α levels in PEX1 deficient cells. Lastly, we demonstrate that upon hypoxia-reoxygenation, PEX1 deficient cells are slower to reset HIF1α levels. Our results suggest that PBD patients with PEX1 deficiency may be susceptible to dysregulation of the HIF1α pathway, particularly in tissues where oxygen gradients are physiological or developmentally required.

## Introduction

The peroxisome is an oxidative organelle that compartmentalizes enzymes that consume oxygen and generate hydrogen peroxide, such as acetyl-CoA oxidase, as well as enzymes that reduce the reactive oxygen byproducts, such as catalase and peroxiredoxins (1, 2). This enables cells to take advantage of oxygen as a terminal electron acceptor, while protecting cells from damage caused by reactive oxygen species. In mammalian cells, peroxisomes coordinate the β-oxidation of very long chain fatty acids and are the sole organelles that perform α-oxidation of fatty acids (3–5). They also contribute to plasmalogen biosynthesis, amino acid metabolism, and glyoxylate metabolism. Peroxisome metabolism is intricately linked to metabolic processes in other organelles, including the ER, lipid droplets, and the mitochondria, which can form contact sites with peroxisomes to exchange lipids, metabolites, and reactive oxygen species (6, 7).

Peroxisomes in various cell types are remodeled according to their specific metabolic needs. The number of peroxisomes in a cell is determined by the balance of *de novo* biogenesis, proliferation by fission, and degradation by peroxisome-specific autophagy (pexophagy) (8, 9). Formation and maintenance of peroxisomes in mammalian cells requires a set of proteins encoded by *PEX* genes, which when mutated, lead to neurodevelopmental disorders known as Peroxisome Biogenesis Disorders (PBDs) (10). PBDs present a wide range of symptom severity. These disorders are characterized by widespread metabolic dysfunction, elevated levels of circulating very long chain fatty acids, and, in the most severe cases, are lethal in early childhood (11). Particular defects associated with peroxisome dysfunction are skeletal dysplasia, chondrodysplasia, hepatomegaly, hepatic insufficiency, as well as nervous system defects such as loss of myelination (12, 13). Milder PBDs manifest as progressive sensorineural hearing loss and retinal degeneration (14, 15).

Most diagnosed PBDs result from mutations in PEX1 which, together with its partner protein PEX6, form a hetero-hexameric Type II AAA-ATPase, that facilitates protein import into the peroxisome lumen (16). The majority of peroxisomal enzymes are targeted to the organelle by a C-terminal tripeptide motif (PTS1) that is recognized by the cytosolic receptor PEX5. During the import process, PEX5 and the PTS1-protein both enter the peroxisome lumen. The PEX1/PEX6 motor is critical for extracting PEX5 from the peroxisome lumen back to the cytosol (17, 18) for subsequent rounds of import. In the absence of PEX1/PEX6 activity, PEX5 accumulates at the peroxisome membrane, where it is poly-ubiquitinated and targeted for degradation (19, 20). PEX1 is also critical for preventing the autophagic degradation of functional peroxisomes by inhibiting NBR1-dependent pexophagy (21, 22), and therefore deleterious mutations in PEX1 or PEX6 result in abrogated matrix protein import, increased peroxisome-specific autophagy, and general peroxisome dysfunction (23).

Previous work has demonstrated that certain metabolic disorders arising due to mitochondrial dysfunction can be exacerbated or alleviated by changes in oxygen pressure. For example, in mice, the symptoms of Leigh syndrome and Friedreich’s ataxia can be alleviated by hypoxia (24, 25). One way mammalian cells react to hypoxia is via the well-studied HIF signaling pathway. The HIF family transcription factors and their regulatory partners form a highly conserved oxygen sensing module in metazoans that allows cells to respond to low oxygen environments by broadly rewiring their metabolism and driving angiogenic programs (26, 27). The HIF transcription factors are heterodimers comprised of a constitutively expressed HIF-1β subunit, and the oxygen regulated HIF1α and HIF2α subunits. In normal oxygen tensions, the HIFα subunits are efficiently hydroxylated by proline hydroxylase domain (PHD) enzymes, and rapidly targeted for proteasomal degradation by a complex containing the E3 ubiquitin ligase von Hippel-Lindau (VHL) tumor suppressor protein (28). In hypoxic conditions, the oxygen-dependent PHD enzymes cease working, stabilizing the HIFα subunits and promoting their dimerization and transcription factor activity (29). These transcription factors therefore reprogram carbohydrate and lipid metabolism along with oxidative stress responsive genes in order to allow cells to survive oxygen depletion (30–32). Upon reoxygenation, PHD enzyme function is restored, leading to rapid degradation of the accumulated HIFα subunits and termination of hypoxic signaling.

While the links between mitochondrial function and HIF activation are well documented, the links between these pathways and peroxisomes are comparatively underexplored. Additionally, there has been no documented testing of whether hypoxia or hyperoxia alters symptoms in models for Peroxisome Biogenesis Disorders. However, the link between peroxisomes and oxygen sensing are being increasingly appreciated: a genome wide screen for genes that confer advantages at different oxygen pressures found that cells with disruptions in peroxisome genes are particularly sensitive to hypoxic conditions. This sensitivity was determined to be due to the toxicity of saturated lipids in hypoxia which can be ameliorated by ether lipid synthesis in the peroxisome (33). It has also been reported that HIF2α induces peroxisome-specific autophagy via NBR1, suggesting that peroxisome degradation is a specific response to hypoxia (34). Further complicating the picture, another group reported that in rodent hepatocytes that have undergone hypoxia followed by reoxygenation, HIF regulatory factors such as the PHD enzymes and VHL localize to the peroxisome (35). This suggests a more complex relationship between oxygen-sensing pathways and peroxisome function than was previously appreciated, one in which remodeling of peroxisome function may be necessary for particular oxygenation states, or upon cycles of cellular hypoxia-reoxygenation.

Here we use a CRISPR-Cas9 edited PEX1^G843D^ cell line and PEX1 null PBD patient fibroblasts to show that PEX1 deficiency results in aberrantly elevated levels of HIF1α protein and transcriptional activity in normoxic conditions. This increase was restored by inducible overexpression of PEX1^WT^, confirming that the effect was dependent on peroxisome function. The protein level increase was not explained by concomitant increases in the *HIF1A* mRNA, or dysfunction of the canonical PHD-VHL-proteasome degradation axis, nor by alterations in global or stress responsive translation. Instead, we show that the difference in HIF1α levels between PEX1 deficient and wildtype cells arises due to impaired autophagy of hydroxylated HIF1α in PEX1 deficient cell lines. The increased level of HIF1α in PEX1 deficient cells could be rescued by increasing autophagic capacity through treatment with an agonist of Unc-51 like autophagy activating kinase 1 (ULK1). While PEX1 deficient cells still react to hypoxia, they are slower to reset HIF1α levels to baseline following hypoxia-re-oxygenation, a process in which a large pool of hydroxylated HIF1α must be rapidly cleared from the cell.

Taken together, we propose a model in which HIF1α can be degraded through a hydroxylation-dependent, autophagic mechanism that is limiting in PEX1 deficient cells. These results corroborate similar findings from other groups that have recently showed impaired autophagic clearance of various substrates in PEX1 deficiency due to an overcommitment of the machinery to pexophagy alone (36). Our results extend these observations to a key pathway of HIF1α degradation and suggest that patients with mutations in PEX1 might be prone to dysregulation of HIF1α signaling in tissues that rely on dynamic responses to physiological oxygen gradients, or in transient hypoxia such as in conditions of ischemia-reperfusion or general anesthesia (37, 38).

## Results

### PEX1 defective cell lines have higher HIF1α protein levels at steady state and show an elevated HIF1α transcriptional activity

We recently demonstrated that the PEX1^G843D^ protein is rapidly degraded by the proteasome, but can form a functional PEX1/PEX6 hetero-hexameric AAA-ATPase complex and support peroxisome function when protected from proteasomal degradation (39). While performing cycloheximide chase assays with proteasome inhibitors, we noticed that HIF1α protein levels were significantly elevated in CRISPR-Cas9 edited PEX1^G843D^ cells compared to PEX1^WT^ HCT116 cells in which HIF1α was hardly detectable at steady state (Fig 1A, B). We then assessed if the levels of HIF1α protein were also elevated in skin fibroblasts from PEX1^-/-^ (null) PBD patients compared to control fibroblasts and again observed a marked increase in total HIF1α (Fig 1C, D). In both cell types the level of Pro564-hydroxylated form of HIF1α was increased in addition to the total level of HIF1α, indicating that proline hydroxylation by PHD is still functional in PEX1 deficient cells.

**Figure 1.**
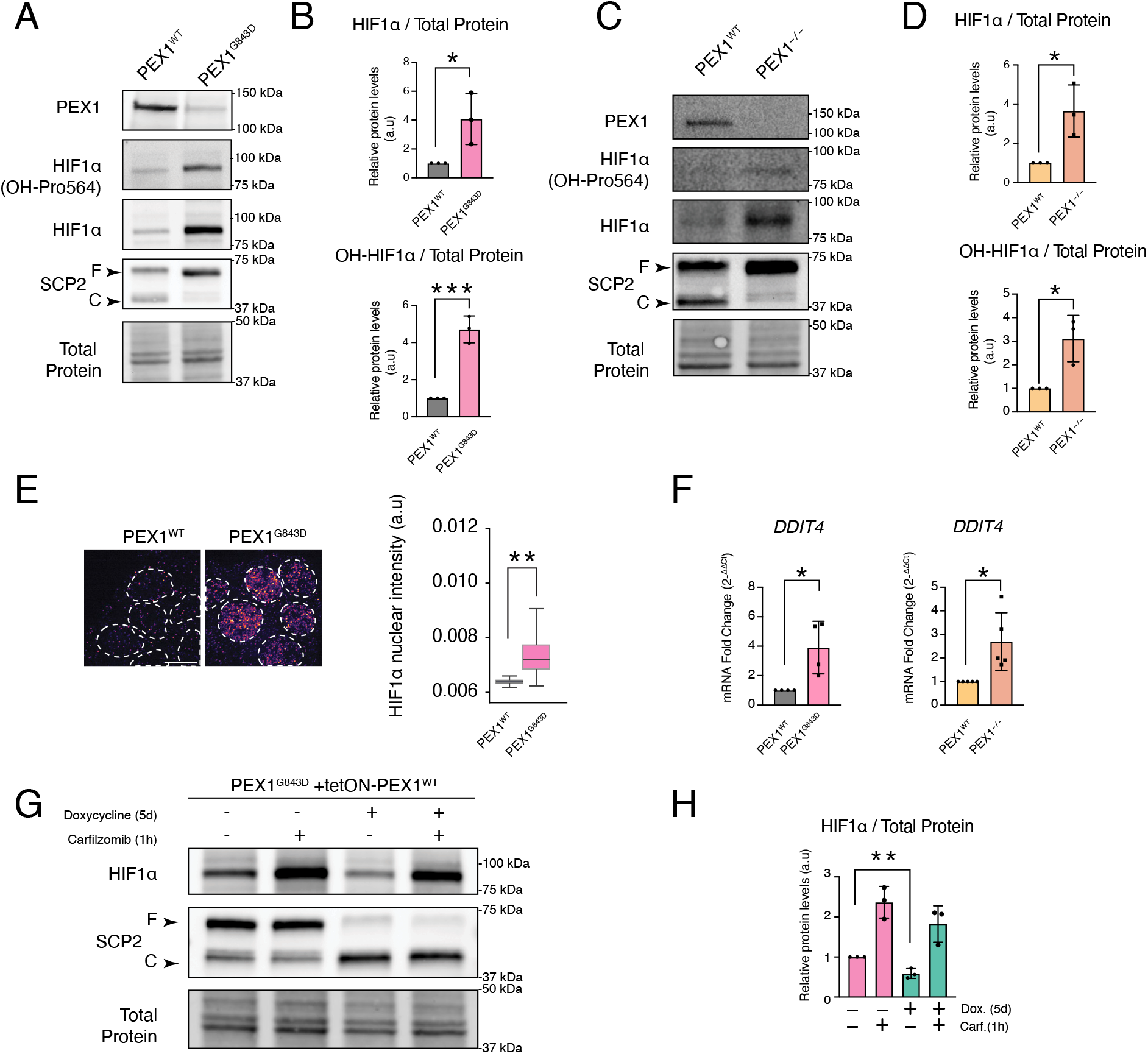
(A) Immunoblot of total and Pro564 hydroxylated HIF1α protein levels in PEX1^WT^ and PEX1^G843D^ HCT116 cells in normoxic conditions. The lack of a cleaved SCP2 band confirms the lack of PTS1 import in PEX1^G843D^ cells (B) Pixel densitometry analysis of total and Pro564 hydroxylated HIF1α levels from *(A).* (C) Immunoblot of total and Pro564 hydroxylated HIF1α protein levels in PEX1^WT^ and PEX1^-/-^ PBD patient fibroblasts in normoxic conditions. The lack of a cleaved SCP2 band confirms the lack of PTS1 import in PEX1^-/-^ fibroblasts (D) Pixel densitometry analysis of total and Pro564 hydroxylated HIF1α levels from *(C)*. (E) Immunofluorescence microscopy of HCT116 PEX1^WT^ and PEX1^G843D^ cells showing nuclear localization of HIF1α at steady state and quantification of per nucleus intensity of HIF1α. The scale bar represents 10µm. (F) ΔΔCt fold change analysis of *DDIT4* mRNA levels by qRT-PCR in PEX1^WT^ and PEX1^G843D^ HCT116 cells, or PEX1^WT^ and PEX1^-/-^ PBD patient derived fibroblasts. (G) Immunoblot of HIF1α in PEX1^G843D^ cells expressing a doxycycline inducible tetON-PEX1^WT^ construct treated with doxycycline for 5d or carfilzomib for 1h or both. Restoration of SCP2 processing in doxycycline treatment confirms rescue of peroxisome function. (H) Pixel densitometry analysis of HIF1α levels in *(G)* Pairwise comparisons were performed using a two-tailed unpaired Student’s t-test, where p < 0.05*, p < 0.01**, p < 0.001***

To assess whether the stabilized and hydroxylated HIF1α protein could be functional within cells, we performed immunofluorescence microscopy for the HIF1α protein in PEX1^WT^ or PEX1^G843D^ HCT116 cells. We observed higher basal levels of HIF1α in the nucleus of PEX1^G843D^ cells compared to PEX1^WT^, suggesting that the stabilized protein was functionally active (Fig 1E). Additionally, we re-analyzed an existing RNA-Seq dataset from our lab comparing HCT116 CRISPRi cells expressing a non-targeting control small guide RNA (sgRNA) to cells expressing a sgRNA targeting PEX1. Differential gene expression analysis confirmed that PEX1 was strongly depleted in the PEX1 sgRNA expressing cells (log2FC = -2.43, adjusted p-value = 1.04E-13). Next, we performed gene set enrichment analysis (GSEA) to assess the effect of PEX1 depletion on HIF1α target genes using published gene sets representing known transcriptional targets of HIF1α (40–42). We found a significant enrichment of these HIF1α targets in PEX1 depleted cells, suggesting that these cells may be experiencing aberrant activation of HIF signaling in normoxic culture conditions. In addition, hallmark gene sets related to reactive oxygen species pathways were significantly enriched in PEX1 depleted cells (FDR ≤ 0.1, NES ≥ 1.5) (Fig S1A). We then performed quantitative RT-PCR for a direct transcriptional target of HIF1α, *DDIT4*, in edited HCT116 PEX1^G843D^ cells as well as PEX1^-/-^ PBD patient fibroblasts and compared their *DDIT4* mRNA levels to matched PEX1^WT^ controls. We found that *DDIT4* expression was enhanced in both models of PEX1 deficiency compared to wildtype cells, confirming low level HIF1α transcriptional activity in normoxic PEX1 deficient cells (Fig 1F).

Next, we tested if the observed elevation in HIF1α protein levels was truly attributable to peroxisome function. We re-expressed PEX1^WT^ from a stably integrated doxycycline inducible promoter in our PEX1^G843D^ edited HCT116 cell line for 5 days. Treatment with doxycycline for 5 days entirely rescued the proteolytic processing of SCP2 within the peroxisome to its lower molecular weight mature form, indicating restored PTS1 import and peroxisome function (Fig 1G). Additionally, the levels of HIF1α were significantly reduced upon doxycycline treatment to restore PEX1 expression, suggesting that proper peroxisome function does indeed lower its accumulation (Fig 1H). Notably, treatment with the irreversible proteasome inhibitor, carfilzomib, increased HIF1α levels in the PEX1^G843D^ cell line with and without the PEX1 addback, a preliminary indication that HIF1α can still be degraded by the proteasome in both conditions (Fig 1G,H).

### PHD-mediated proline hydroxylation, VHL-mediated ubiquitination, and proteasomal degradation remain active in PEX1 deficient cell lines

As an oxygen sensitive transcription factor, HIF1α stabilization is tightly coupled to oxygen tension. In oxygen replete conditions, the HIF1α protein is rapidly hydroxylated by the oxygen-dependent PHD1/2/3 family of prolyl hydroxylase enzymes, subsequently ubiquitinated by the E3 ubiquitin ligase VHL, and targeted for proteasomal degradation or autophagic degradation through separate recognition and targeting machinery (Fig 2A). We first assessed if the increased levels of HIF1α were correlated with mRNA level changes in either *HIF1A* or *VHL*. To test this, we performed qRT-PCR for the two gene transcripts in PEX1^WT^ and PEX1^-/-^ PBD fibroblasts, as well as HCT116 PEX1^WT^ and PEX1^G843D^ cells. We found that the mRNA transcript levels of the *HIF1A* and *VHL* genes were not significantly different between PEX1^WT^ and PEX1^-/-^ patient fibroblasts as well as between PEX1^WT^ and PEX1^G843D^ cells, suggesting that mRNA level regulation was insufficient to explain our observations (Fig 2B, C).

**Figure 2.**
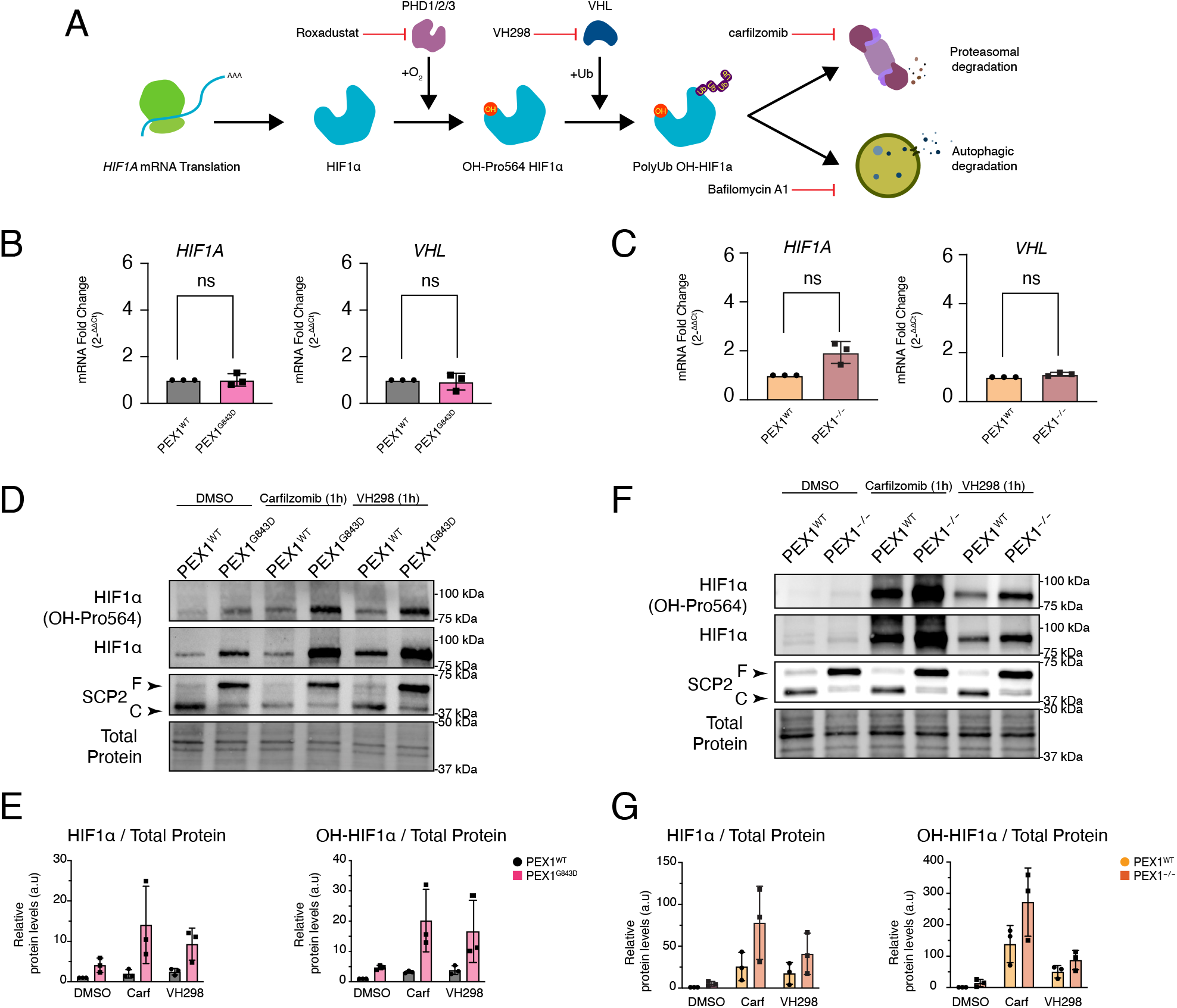
(A) Schematic of HIF1α mapping its fate from translation to degradation, highlighting its regulation by oxygen dependent proline hydroxylases, PHD1-3, and the E3 ubiquitin ligase VHL that targets it for proteasomal degradation as well as pharmacological inhibitors used in the remainder of this study, and their respective mechanisms of action (B) ΔΔCt fold change analysis of *HIF1A* and *VHL* mRNA transcripts by qRT-PCR in HCT116 PEX1^WT^ or PEX1^G843D^ cells. (C) ΔΔCt fold change analysis of *HIF1A* and *VHL* mRNA transcripts by qRT-PCR in PEX1^WT^ and PEX1^-/-^ PBD patient fibroblasts. (D) Immunoblot of PEX1^WT^ or PEX1^G843D^ HCT116 cells treated with DMSO, carfilzomib, or VH298 for 1h showing the levels of total and Pro564 hydroxylated HIF1α, as well as SCP2 processing as a proxy for peroxisome import (E) Pixel densitometry analysis of total and Pro564 hydroxylated HIF1α from *(D).* (F) Immunoblot of PEX1^WT^ or PEX1^-/-^ PBD patient fibroblasts treated with DMSO, carfilzomib, or VH298 for 1h showing the levels of total and Pro564 hydroxylated HIF1α, as well as SCP2 processing as a proxy for peroxisome import. (G) Pixel densitometry analysis of total and Pro564 hydroxylated HIF1α from *(F)* Pairwise comparisons were performed using a two-tailed unpaired Student’s t-test, where p < 0.05*, p < 0.01**, p < 0.001***

Our initial observations indicated that PHD-dependent proline hydroxylation at the residue known to be required for oxygen dependent HIF1α degradation remained robust in PEX1 deficient cells (Fig 1A, B, C, D); as we detected increased levels of both total and Pro564-hydroxylated HIF1α. Additionally, proteasomal degradation of HIF1α also appeared robust in PEX1 deficient cells (Fig 1G). To further dissect any alterations of the canonical PHD-VHL degradation pathway, we treated PEX1^WT^ and PEX1^G843D^ HCT116 cells with either carfilzomib (an irreversible proteasome inhibitor), or VH298 (an inhibitor of the VHL E3 ubiquitin ligase) and performed immunoblot analysis. With both inhibitors, the level of total and hydroxylated HIF1α in PEX1 deficient cells was elevated compared to the DMSO treatment conditions (Fig 2D, E). These results indicate that PEX1 deficient cells are able to degrade normoxic levels of HIF1α robustly through VHL ubiquitination and proteasomal degradation. Furthermore, in both treatment conditions, there was more HIF1α in PEX1-deficient cell lines than in WT, indicating that the disparity in HIF1α levels does not arise through reduced VHL-ubiquitination and proteasomal degradation in PEX1 deficient cell lines.

We performed the same experiment in PEX1^WT^ and PEX1^-/-^ PBD patient fibroblasts and observed a similar pattern: PEX1^-/-^ cells accumulated the HIF1α protein to a much greater degree than PEX1^WT^ upon inhibition of VHL-proteasome dependent degradation (Fig 2F, G). These results suggest that HIF1α ubiquitination and proteasomal degradation is active in PEX1 deficient cell lines and that dysfunction of this pathway is insufficient to explain the elevation in protein levels in normoxic conditions. Given our results with proteasome and VHL inhibitors, and the well-known regulatory mechanisms that control HIF1α expression at the translational level, we wondered whether the observed increase in protein levels was explainable by alterations in global or stress responsive translational control.

HIF1α translation can be controlled by an internal ribosome entry site that may allow for stress dependent translation in contexts where global cap dependent translation is inhibited, such as in conditions of oxidative or nutrient stress (43). These diverse cellular stressors are detected by the four sensor kinases of the Integrated Stress Response which converge on the phosphorylation of the α subunit of the eukaryotic initiation factor eIF2, reducing the availability of the ternary complex required for cap-dependent translational initiation (44). However, when we analyzed the levels of phosphorylated and total eIF2α by immunoblot, we did not observe any significant differences between PEX1^WT^ and PEX1^G843D^ cells (Fig S2A, B) suggesting that stress responsive translation is insufficient to explain our observations. To further corroborate these observations, we assessed the levels of global translation in our cells by performing a puromycylation pulse labeling experiment where we treated PEX1^WT^ or PEX1^G843D^ cells with either DMSO or cycloheximide for 1 hour, followed by puromycin labeling for 10 minutes. We did not observe an increase in global cap-dependent translation, and in fact observed a slight decrease in PEX1^G843D^ cells instead, suggesting that global translational activation was not the cause of elevated HIF1α levels (Fig S2C, D)

HIF1α translation is also known to be positively regulated by the activation of mTORC1 (45). Thus, we assessed the activation status of mTORC1 by immunoblot analysis of its direct phosphorylation target, P70S6K, as well as the phosphorylation target of P70S6K itself, ribosomal protein S6 (RS6). We did not observe activation of the mTORC1 pathway, and in fact observed a reduced proportion of phosphorylated P70S6K and RS6 in PEX1^G843D^ cells, indicating that PEX1 deficiencies result in a dampened mTORC1 signaling state (Fig S2E, F, G)

### A hydroxylation-dependent, proteasome-independent HIF1α degradation route is defective in PEX1 deficient cells

To further assess flux through the ubiquitin proteasome system in cells lacking PEX1, we performed a cycloheximide washout assay using PEX1^WT^ or PEX1^G843D^ to assess the kinetics of HIF1α degradation and production in each cell line. In this experiment, we first treated the cells with cycloheximide (an inhibitor of translation elongation) for 1h to allow the short-lived HIF1α protein to be turned over. Then, the cycloheximide was washed away, allowing translation to resume, but the cells were recovered in the presence of VH298 to inhibit VHL dependent ubiquitination. Therefore, the cells were allowed to resume translation but were prevented from degrading newly made HIF1α through the VHL-proteasome route. Prior to cycloheximide treatment, HIF1α levels were elevated in PEX1^G843D^ cells consistent with our prior experiments. After 1h of cycloheximide treatment, both cell lines had degraded their pool of HIF1α to barely detectable levels confirming that oxygen dependent degradation pathways remained active in PEX1^G843D^ cells. When translation was resumed in the presence of VH298, PEX1^G843D^ cells quickly accumulated large amounts of total and Pro564 hydroxylated forms of HIF1α, whereas the HIF1α levels in PEX1^WT^ cells increased only slightly (Fig 3A,B). We also performed this experiment using wildtype or PEX1^-/-^ PBD patient fibroblasts and observed similar results where the cells lacking PEX1 accumulated HIF1α to a greater degree (Fig S3A, S3B). These results indicate that the difference arising between PEX1^WT^ and PEX1 deficient cell lines is not dependent on VHL-mediated ubiquitination. Having already ruled out increased translation of HIF1α, we hypothesized that a VHL independent degradation pathway allows HIF1α clearance in PEX1^WT^ cells, and that this pathway is dysfunctional in PEX1 deficient cells.

**Figure 3.**
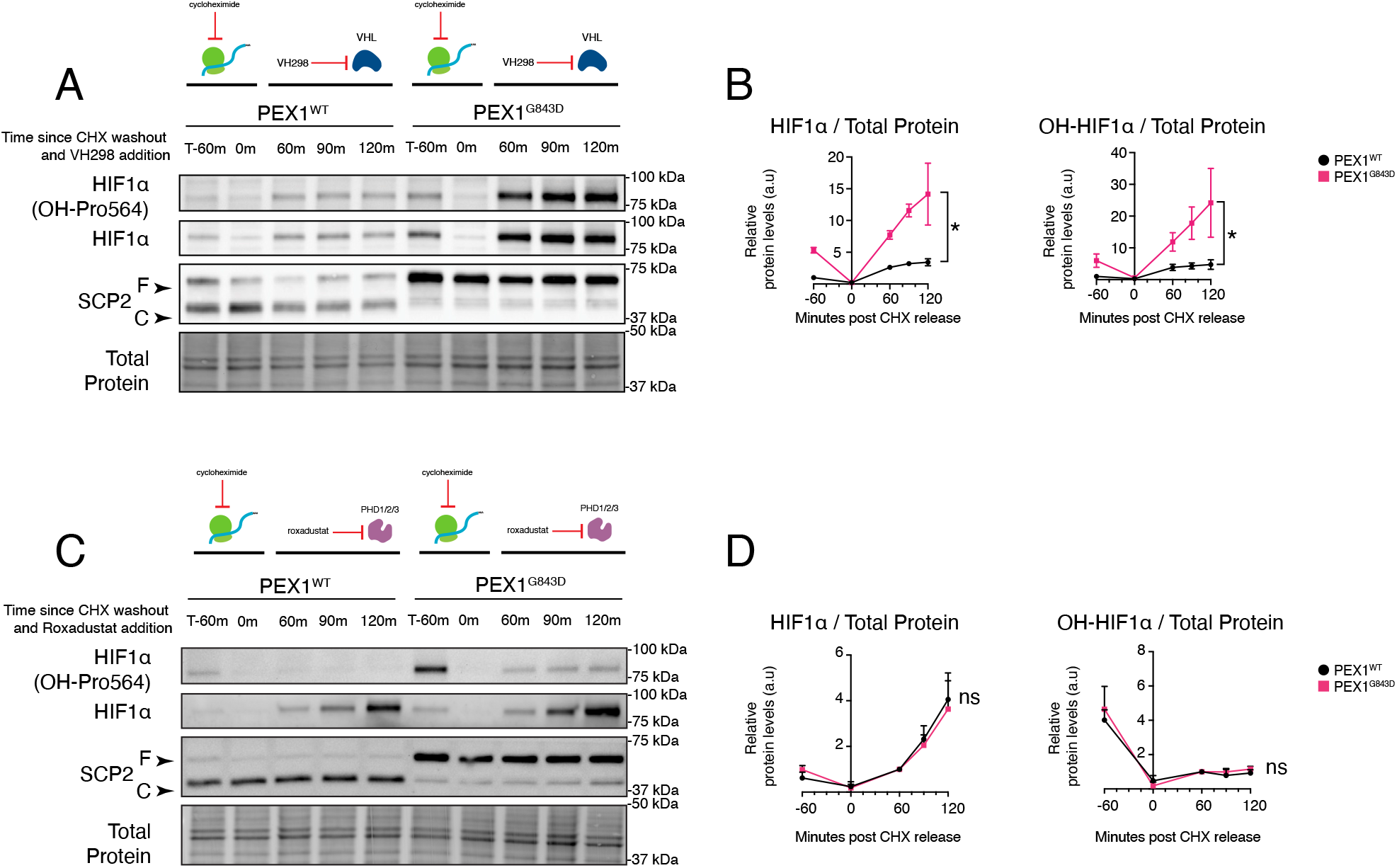
(A) Immunoblot of total and Pro564 hydroxylated HIF1α in PEX1^WT^ and PEX1^G843D^ cells in a cycloheximide (CHX) washout experiment. Cells are first treated with CHX for 1h to allow complete degradation of HIF1α to almost zero. Then CHX is washed out and replaced with medium containing VH298 to assess the build-up of HIF1α over time when VHL dependent degradation is blocked. The SCP2 immunoblot confirms a lack of peroxisomal import in the PEX1^G843D^ cell line. (B) Pixel densitometry analysis of total HIF1α levels over time from *(A).* (C) Immunoblot of total and Pro564 hydroxylated HIF1α in PEX1^WT^ and PEX1^G843D^ cells in a cycloheximide (CHX) washout experiment. Cells are first treated with CHX for 1h to allow complete degradation of HIF1α to almost zero. Then CHX is washed out and replaced with medium containing Roxadustat to assess the build-up of HIF1α over time when PHD dependent hydroxylation is blocked. The SCP2 immunoblot confirms a lack of peroxisomal import in the PEX1^G843D^ cell line. (D) Pixel densitometry analysis of total and Pro564 hydroxylated HIF1α levels over time from *(C).* Pairwise comparisons were performed using a two-tailed unpaired Student’s t-test, where p < 0.05*, p < 0.01**, p < 0.001***

To test if the discrepancy in HIF1α depends on proline hydroxylation, we repeated the cycloheximide washout assay allowing the cells to resume translation in the presence of an inhibitor of the prolyl hydroxylase enzyme family, Roxadustat, rather than VH298. Surprisingly, when cells resumed translation in the presence of Roxadustat, the accumulation of total HIF1α in PEX1^WT^ cells was equivalent to the levels in PEX1^G843D^ cells (Fig 3C, D). The levels of Pro564 hydroxylated HIF1α did not increase after resumption of translation, confirming that Roxadustat was effective in inhibiting the PHD hydroxylases (Fig 3D). Given the difference in phenotype between VH298 and Roxadustat treatment, the difference arising between PEX1^WT^ and PEX1 deficient cell lines is dependent on PHD-dependent hydroxylation but not VHL-ubiquitination or proteasomal degradation. We surmise that wildtype cells are able to degrade HIF1α through a hydroxylation-dependent, but VHL- and proteasome-independent mechanism, and that this mechanism is defective or limiting in cells lacking PEX1.

### Autophagic clearance of hydroxylated HIF1α is limiting in PEX1 deficient cells, but rescued by increasing autophagic capacity via ULK1 agonism

The degradation of HIF1α through an autophagy dependent mechanism has been reported by several groups (46, 47). Therefore, we wondered if the hydroxylation-dependent, but VHL- and proteasome-independent mechanism of HIF1α degradation in PEX1^WT^ cells depends on the availability of autophagic machinery. We thus repeated our cycloheximide washout assay but allowed cells to resume translation in the presence of VH298 (an inhibitor of the VHL E3 ubiquitin ligase) as well as an inhibitor of autophagosome-lysosome fusion, Bafilomycin A1. Here, we observed that PEX1^WT^ cells now matched the rate of accumulation observed in PEX1 deficient cells (Fig 4A). Therefore, PEX1^WT^ cells accumulate significantly more HIF1α protein when both VHL ubiquitination and autophagy are inhibited compared to when only VHL ubiquitination is inhibited. The Pro564 hydroxylated version of the HIF1α protein also accumulated to the same extent in both cell lines, indicating that PHD hydroxylation remains active (Fig 4A, B). Together, these results suggest that autophagic clearance of hydroxylated HIF1α is impaired in PEX1 deficient cell lines, which leads to an increase in HIF1α levels.

**Figure 4.**
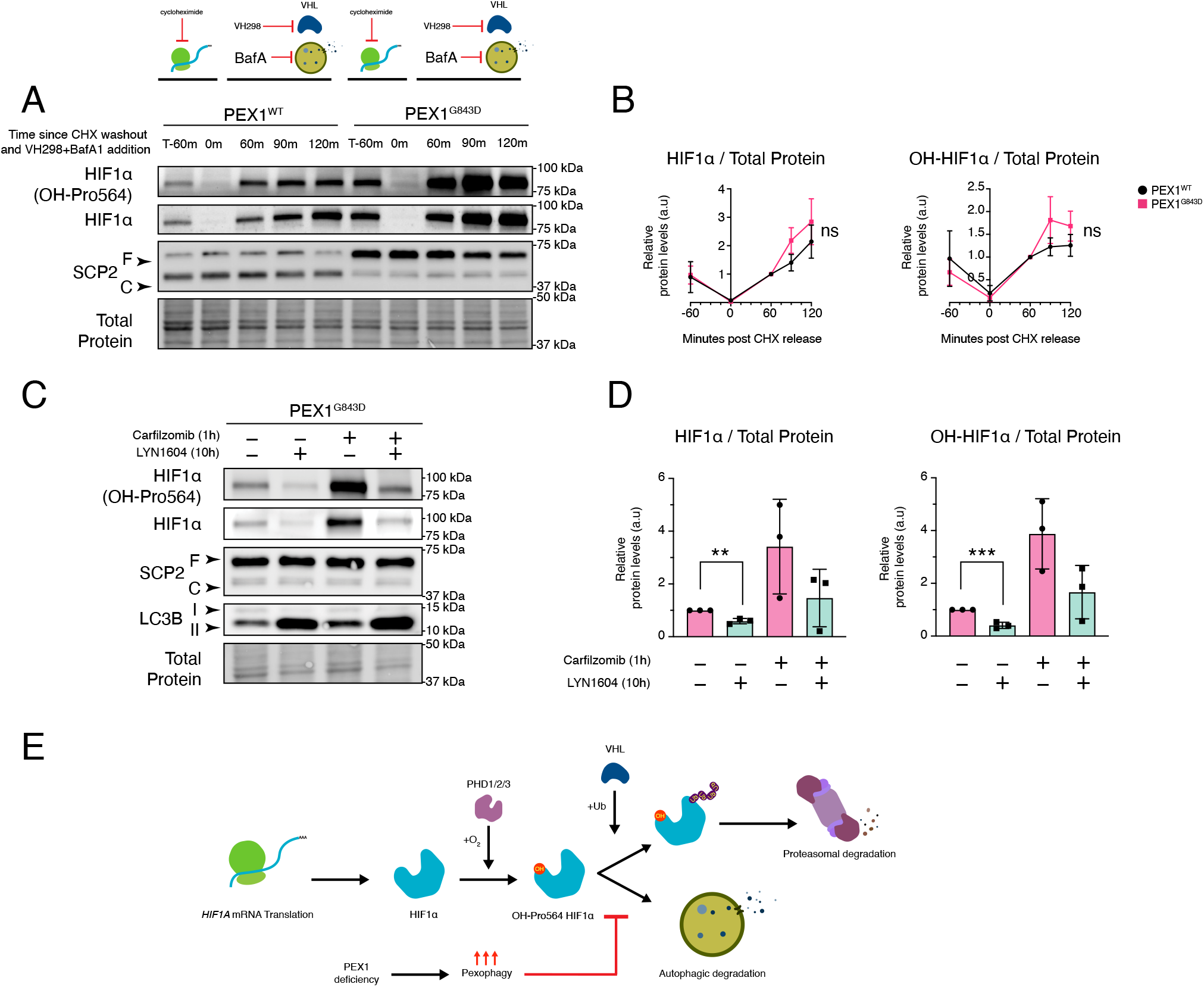
(A) Immunoblot of total and Pro564 hydroxylated HIF1α in PEX1^WT^ and PEX1^G843D^ cells in a cycloheximide (CHX) washout experiment. Cells are first treated with CHX for 1h to allow complete degradation of HIF1α to almost zero. Then CHX is washed out and replaced with medium containing VH298 and Bafilomycin A1 to assess the build-up of HIF1α over time when VHL dependent and autophagy dependent degradation are both blocked. The SCP2 immunoblot confirms a lack of peroxisomal import in the PEX1^G843D^ cell line. (B) Pixel densitometry analysis of total and Pro564 hydroxylated HIF1α levels over time from *(A)* (C) Immunoblot of PEX1^G843D^ cells treated for 10h with the ULK1 agonist LYN-1604 prior to treatment with DMSO or Carfilzomib for 1h. The LC3B immunoblot indicates sufficiently increased autophagic flux by the lower ratio of LC3BI/LC3BII (D) Pixel densitometry analysis of total and Pro564 hydroxylated HIF1α from *(C)* (E) Model of HIF1α degradation through a parallel hydroxylation and autophagy dependent mechanism that is compromised in PEX1 deficiency. Pairwise comparisons were performed using a two-tailed unpaired Student’s t-test, where p < 0.05*, p < 0.01**, p < 0.001***

It has previously been shown that PEX1 deficiencies, including PEX1^G843D^, trigger peroxisome specific autophagy (21). This results in an overcommitment of the autophagic machinery, compromising the clearance of other substrates (36). Our result suggests that HIF1α may be one such client that aberrantly accumulates in PEX1 deficiencies due to saturated autophagic capacity. To test this model, we treated PEX1^G843D^ cells with LYN-1604, an agonist of the autophagy initiator, ULK1, that broadly increases the capacity of cells to initiate autophagosome nucleation (48, 49). Treatment of LYN-1604 alone or LYN-1604 in combination with carfilzomib reduced the levels of both total and Pro564 hydroxylated HIF1α in PEX1^G843D^ cells, indicating that increased autophagic flux was sufficient to restore its alternative autophagic degradation route, even when the proteasomal degradation route was rendered non-functional (Fig 4C, D). Therefore, our data suggest a model in which hydroxylated HIF1α is constitutively degraded by two partially redundant routes, and that the autophagic route is impaired in PEX1 deficient cells. Our results thus support the model that PEX1 deficiency results in impaired autophagy through chronically elevated pexophagy that sequesters autophagic capacity away from other clients such as HIF1α (Fig 4E).

### PEX1 deficient cells respond like wildtype cells to hypoxia or other hydroxylation inhibitors, but are slower to reset HIF1α to baseline upon hypoxia-reoxygenation

What are the consequences of impaired autophagic degradation of HIF1α in PEX1 deficient cells? Since the observed difference in HIF1α degradation depends on oxygen-dependent hydroxylation, we reasoned that the difference between WT and PEX1 deficient cells would be reduced in conditions that are non-permissive to hydroxylation. Therefore, treatment with compounds such as Roxadustat or CoCl_2_, (both of which mimic hypoxia by inhibiting the prolyl hydroxylase domain family of enzymes) or exposure to true hypoxia should reduce any differences between wildtype cells and cells lacking PEX1. To test this, we treated PEX1^WT^ or PEX1^G843D^ cells for 4 hours with CoCl_2_ or exposed them to 1% O_2_ for 4 hours. We observed that the normoxic condition still showed elevated levels of HIF1α in PEX1 deficient cells compared to wildtype, but treatment with CoCl_2_ or hypoxia largely abolished these differences (Fig 5A,B). In addition, HIF1α and HIF2α accumulation were matched in both cell lines indicating that the ability to mount a hypoxic response was not compromised in PEX1 deficient cells (Fig 5A, B). To confirm this observation, we performed the same experiment but measured HIF1α nuclear intensity by immunofluorescence staining instead. Similar to our results with the immunoblot experiment, normoxic PEX1^G843D^ cells showed increased levels of nuclear HIF1α signal compared to PEX1^WT^, but upon treatment with hypoxia or CoCl_2_ both cell lines accumulated nuclear HIF1α maximally and to similar levels (Fig 5C, D).

**Figure 5.**
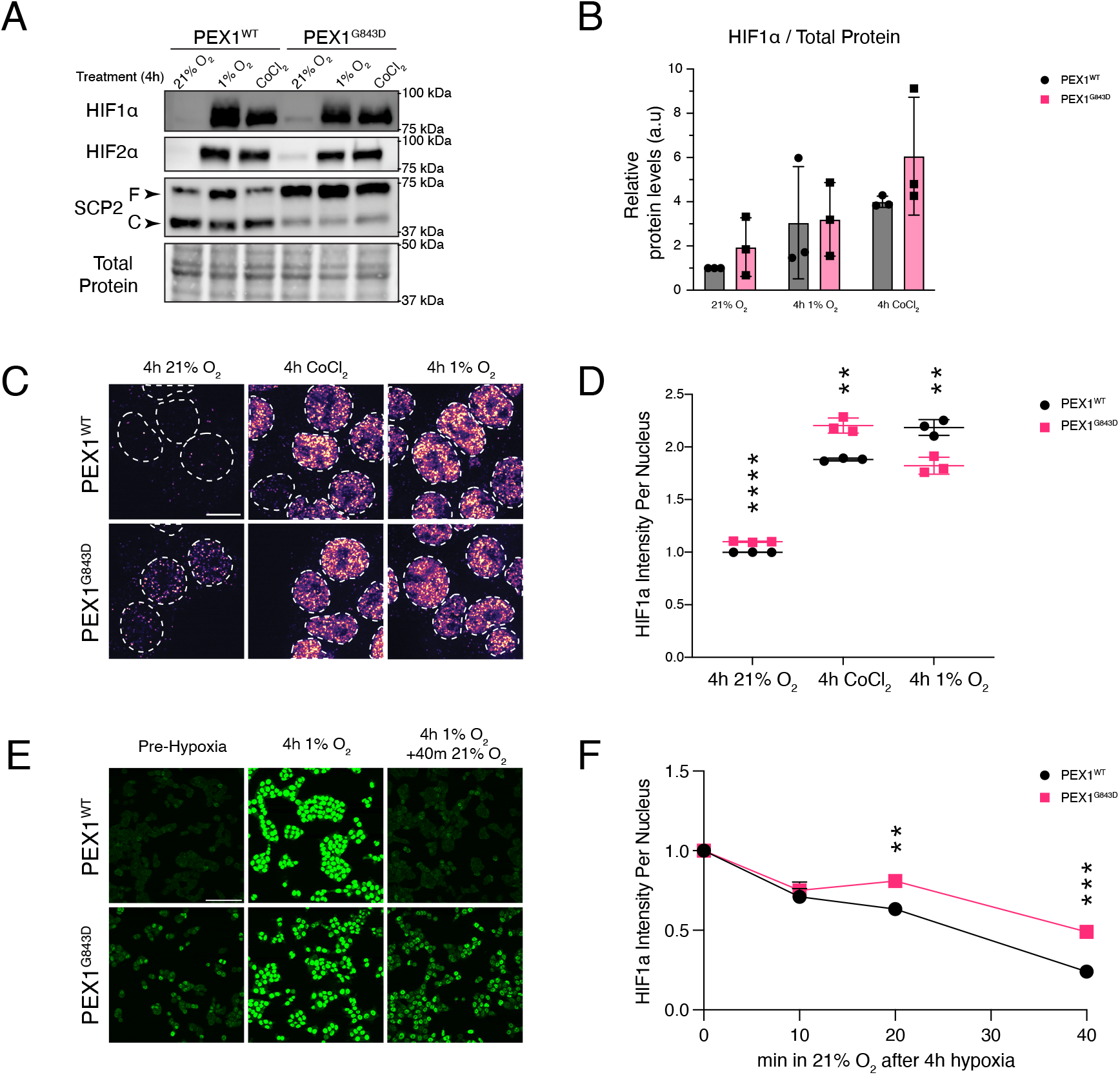
(A) Immunoblot of total and Pro564 hydroxylated HIF1α and total HIF2α in PEX1^WT^ and PEX1^G843D^ cells subjected to 1% O_2_ or CoCl_2_ treatment for 4h. The SCP2 immunoblot confirms a lack of peroxisomal import in the PEX1^G843D^ cell line. (B) Pixel densitometry analysis of total HIF1α from *(A)* (C) Immunofluorescence microscopy of HCT116 PEX1^WT^ and PEX1^G843D^ cells showing nuclear localization of HIF1α at steady state and accumulation upon 1% O_2_ or CoCl_2_ treatment (D) Per nucleus intensity of HIF1α quantified from *(C).* The scale bar represents 10µm (E) Immunofluorescence microscopy of HCT116 PEX1^WT^ and PEX1^G843D^ cells showing nuclear localization of HIF1α at steady state, accumulation upon 1% O_2_ treatment for 4 hours, and after 40 minutes of reoxygenation (F) Per nucleus intensity of HIF1α over time quantified from *(E)*. The scale bar represents 100µm. Pairwise comparisons were performed using a two-tailed unpaired Student’s t-test, where p < 0.05*, p < 0.01**, p < 0.001***

Given that PEX1 deficient cells appeared to be able to mount a hypoxic response similar in magnitude to wildtype cells, we hypothesized that the hydroxylation and autophagy dependent degradation route would be crucial in conditions of hypoxia-reoxygenation. In this scenario, HIF1α protein accumulates maximally as hydroxylation by the oxygen-dependent PHD enzymes is inhibited, allowing the cells to initiate a hypoxic response. Upon reoxygenation, this large pool of protein must be efficiently hydroxylated and rapidly degraded to allow HIF1α signaling to reset to baseline. In these conditions, it is possible that the VHL-proteasome dependent degradation route becomes rate-limiting and bulk autophagic degradation is more efficient. Therefore, our data predict that PEX1 deficient cells will experience prolonged HIF1α signaling upon reoxygenation as they degrade the large pool of accumulated HIF1α through the proteasomal route alone, while the wildtype cells can accomplish this with both the proteasomal and autophagic routes in parallel. To test our hypothesis, we performed a hypoxia-reoxygenation experiment in which cells were incubated for four hours in 1% O_2_, then recovered into normoxic cell culture conditions, and fixed and immunostained for HIF1α at 10-, 20-, and 40-minutes post re-oxygenation. We found that after 4 hours of hypoxia, both cell lines accumulated HIF1α to similar degrees even though the normoxic baseline was higher in PEX1^G843D^ cells (Fig 5E). Upon reoxygenation, the PEX1^G843D^ cells cleared HIF1α at a reduced rate compared to wildtype. At 40 minutes post re-oxygenation, the PEX1^G843D^ cells retained significant amounts of the protein in the nucleus compared to PEX1^WT^ (Fig 5E, F). Altogether, our results suggest a model in which PEX1-dysfunction significantly impairs the bulk autophagic clearance of hydroxylated HIF1α, resulting in increased baseline levels of hydroxylated HIF1α and sustained signaling in conditions such as hypoxia-reoxygenation where the flux through the VHL-proteasome route is insufficient to efficiently degrade the accumulated pool of protein.

## Discussion

Peroxisome dysfunction impairs specialized metabolism, with noted impacts on mitochondrial function and organization, as well as ribosomal biogenesis (50). Here we show that PEX1 deficiency also alters the cellular response to hypoxia through impaired degradation of the oxygen sensitive transcription factor HIF1α.

Our results show that severe PEX1 deficiency results in a mild HIF1α transcriptional signature and elevated steady-state levels of HIF1α protein under normoxic conditions. We find that this elevation is not attributable to transcriptional activation of the *HIF1A* gene or dysfunction of the canonical oxygen-dependent degradation pathway. Instead, our data support a model in which high levels of peroxisome-specific autophagy caused by PEX1 deficiency impairs the efficient autophagic clearance of hydroxylated HIF1α, thus shifting the degradative burden of HIF1α abnormally onto the VHL-proteasome axis.

The finding that *HIF1A* mRNA levels remain unchanged in both PEX1^G843D^ HCT116 cells and PEX1^-/-^ patient fibroblasts, distinguishes our model from transcriptional activation of the *HIF1A* locus as observed in response to chronic hypoxia or inflammatory signaling (51). It is plausible that peroxisome dysfunction elicits a state of cellular ‘pseudohypoxia’ with increased levels of H_2_O_2_ and ROS, or alters the availability of PHD substrates such as 2-oxoglutarate, or other regulatory cofactors. However, we found that PEX1 deficient cells show higher levels of Pro564-hydroxylated HIF1α indicating that PHD enzyme activity is sufficient to handle the substrate pool available in those cells. VHL-dependent proteasomal targeting is likewise preserved in PEX1 deficient cells, as demonstrated by HIF1α accumulation upon treatment with both carfilzomib and the VHL inhibitor, VH298. Taken together, these results indicate that the oxygen-dependent degradation machinery is functional in PEX1 deficient cells.

HIF1α protein levels are also known to be regulated by direct translational control with contributions from its 5’ untranslated region, from activators of protein synthesis such as mTORC1 signaling, or upregulation of specific translation factors in hypoxic conditions (45). While we cannot rule out all mechanisms of possible translational activation, we found a mild decrease in global translation in PEX1^G843D^ as assessed by puromycylation of nascent peptides. Additionally, we did not observe signs of Integrated Stress Response activation as assessed by the fraction of phosphorylated eIF2α. We also observed a suppressed mTORC1 signaling state, as assessed by phosphorylated fractions of P70S6K and ribosomal protein S6, suggesting that mTORC1 dependent translational activation is unlikely to explain the stabilization we observed. Notably, HIF1α signaling is reciprocal to mTORC1 signaling: the HIF1α transcriptional target DDIT4, also known as REDD1, is a known negative regulator of mTORC1 that acts to promote TSC1/TSC2 activity that restricts mTORC1 activation (52). Therefore, our observation that PEX1 deficient cells express more *DDIT4* mRNA is consistent with the mTORC1 signaling state we observe. Our findings that peroxisome dysfunction results in chronic mTORC1 inhibition are also consistent with reports from other groups studying induced peroxisome dysfunction in human cells (53).

Why might PEX1 deficient cells have reduced autophagy of HIF1α? One possible model is that the autophagic system is capacity-limited as a result of PEX1 loss and peroxisome dysfunction: chronically dysfunctional peroxisomes are themselves continuous pexophagy cargo, and this sustained demand may saturate autophagic throughput in a way that leaves insufficient capacity for non-peroxisomal substrates such as HIF1α. Autophagy does seem to be induced in PEX1^G843D^ cells as assessed by an immunoblot of LC3-I/II processing (54), lending support to this model. Providing further support for this model, we find that treatment with a ULK1 agonist, LYN-1604, increases autophagic capacity as measured by the ratio of LC3-I/II, and also reduces the levels of HIF1α in PEX1^G843D^ cells. Other groups have also shown that increased pexophagy burden can indeed limit autophagic clearance of other substrates that rely on the same machinery due to exhaustion of the canonical autophagosome precursor machinery (36).

These results also inform the interpretation of our PHD inhibition experiments. When hydroxylation is blocked by Roxadustat, hypoxia, or cobalt chloride, the differences in HIF1α levels between PEX1^WT^ and PEX1 deficient cells are reduced. If the autophagic pathway responsible for HIF1α clearance targets the proline-hydroxylated form of the protein, under conditions of PHD inhibition, non-hydroxylated HIF1α accumulates that is recognized neither by VHL nor by the autophagy machinery, causing both routes to fail simultaneously in both cell lines. An implication of this model is that the HIF1α proteostasis deficit in PEX1 deficient cells is fundamentally specific to normoxic conditions, resulting in a higher baseline level of hydroxylated HIF1α. Under severe hypoxia, PHD inhibition by low oxygen would disable autophagic as well as proteasomal recognition of HIF1α in both wildtype and PEX1^G843D^ cells simultaneously, reducing any differences. The relevant phenotype is therefore not exaggerated hypoxic HIF1α induction, but rather impaired normoxic clearance. In conditions of reoxygenation and recovery, the large pool of HIF1α accumulated during hypoxic exposure becomes rapidly hydroxylated and must be efficiently degraded to reset HIF1α signaling to baseline. Therefore, our model predicts that clearance rates upon reoxygenation would be significantly delayed in PEX1 deficient cells with a defective autophagy axis compared to PEX1^WT^ cells. Indeed, when we exposed cells to hypoxic conditions for four hours and then returned them to normoxia, PEX1^WT^ cells rapidly degraded the accumulated HIF1α protein within 40 minutes, while PEX1^G843D^ cells retained significant amounts of nuclear HIF1α signal by this time point. This may become particularly relevant during physiological or developmental oscillations in tissue oxygen tension, or during the recovery phase following transient mild hypoxemic episodes such as in conditions of ischemia-reperfusion or general anesthesia. Interestingly, conditions such as viral infection that result in increased pexophagy (55) are also predicted by HIF1α dynamics models to exhibit deficits in metabolic recovery from hypoxia, and in reduced hysteresis upon recovery (56).

Our findings are intriguing in light of the tissues most severely affected in PBD patients. The liver, skeletal system, and inner ear are all affected in PBD patients, with hepatomegaly, cholestasis, skeletal dysplasia, and sensorineural hearing loss being common symptoms (14). Given that these tissues are ones in which physiological oxygen gradients are functionally required, it is interesting to speculate on how peroxisomes may coordinate cellular processes through HIF signaling. In the liver for example, the periportal-to-pericentral oxygen gradient coordinates zone-specific metabolic programs, where HIF1α and HIF2α are known regulators of lipogenesis related gene expression in the pericentral zone (30). Peroxisome abundance also correlates negatively with oxygen concentration in certain organs with physiological oxygen gradients, such as in the liver where they are more densely populated near the oxygen poor central vein compared to the oxygen rich portal vein (38). In bone marrow, physiological hypoxia directs the spatial coupling of osteogenesis and angiogenesis, with HIF1α activating VEGF and other osteogenic factors in response to the oxygen gradient (57, 58). In the cochlea, transient spikes in HIF1α signaling in response to noise-induced ROS has been implicated in signaling cascades protective against noise-induced hearing loss (59–61). In each of these tissues, the pathological consequence of impaired normoxic HIF1α clearance would not be a failure to activate HIF signaling when needed, but a failure to terminate it, resulting in prolonged HIF1α activity following physiological or stress-induced variation in oxygen availability.

Finally, our findings point toward a potentially reciprocal and self-reinforcing relationship between peroxisome biology and HIF signaling more broadly. HIF2α has been demonstrated to be a negative regulator of peroxisome abundance through NBR1-dependent pexophagy, and peroxisome numbers are reduced in VHL-deficient tumors with correspondingly high HIF2α levels (34). Our data suggest that the converse relationship also exists in which peroxisomes modulate the amplitude of HIF signaling. For example, in hypoxic conditions, the stabilization of HIF2α may result in elevated pexophagy and therefore allow sustained HIF1α signaling by sequestering autophagic machinery. It is interesting to note that other groups have demonstrated that peroxisome loss and the resulting inability to synthesize ether lipids sensitizes cells to low oxygen conditions due to lipid saturation stress. Our results lead us to speculate that coordinated degradation and reformation of peroxisomes is required to re-wire peroxisome metabolism to support survival from hypoxic insults.

In the context of PBDs, these observations raise the possibility of a feed-forward loop in which PEX1 deficiency elevates HIF1α by impairing its normoxic clearance, which in turn drives further pexophagy and peroxisome loss through aberrant low-level hypoxic signaling, amplifying the initial deficit. Whether such a loop operates *in vivo* in PBD patient tissues, and whether it can be interrupted therapeutically, are questions of interest for future research. Strategies that enhance peroxisome biogenesis or that directly augment autophagic clearance of HIF1α may represent avenues for any disease context characterized by age- or stress-related peroxisome decline and subsequent dysregulation of redox homeostasis within the cell.

## Data Availability

Data are available upon request

## Supporting Information

This article contains supporting information

## Acknowledgements

We thank Dr. Nancy Braverman and Dr. Catherine Argyriou for the gift of PBD patient fibroblast cell lines. We thank Dr. Meghan Morrissey for allowing us to utilize the lab’s spinning disc confocal microscope. Brooke Gardner acknowledges support from K99/R00GM121880, R35GM146784, and the Searle Scholars Program. Chris Richardson acknowledges support from R35 GM142975. Soham Chowdhury acknowledges support from the Storke Family Fellowship. Sambhav Jain acknowledges support from T34GM136466. Connor Sheedy acknowledges support from the UC Santa Barbara Chancellor’s Fellowship. We thank Christine Joyce, Elizabeth Sharpe, Maddalena Nano, and Molly Kirk at UCSB for valuable feedback and discussions. We thank the Weimbs lab for use of their hypoxia incubator. We thank the Ma lab at UCSB for sharing their puromycin antibody. The content is solely the responsibility of the authors and does not necessarily represent the official views of the National Institutes of Health.

## Author CRediT Statement

S. P. Chowdhury: Conceptualization, Methodology, Validation, Formal analysis, Investigation, Writing – Original draft, Writing – Review and editing, visualization

S. Jain: Methodology, Validation, Formal analysis, Investigation, Writing – Review and editing, visualization

M. Esmerode: Investigation, Validation, Formal analysis, Writing – Review and editing

H. Dong: Validation, Methodology

J. T. Vu: Investigation

C. J. Sheedy: Validation

C. D. Richardson: Formal analysis, Resources, Data Curation

B. M. Gardner: Conceptualization, Funding acquisition, Project administration, Supervision, Writing – Original Draft, Writing – Review and editing

## Methods and Materials

### Cell culture

HCT116dCas9 cells were a gift from the Corn Lab at UC Berkeley. An N-terminal FLAG tag and G843D mutation were introduced into PEX1 by Cas9 editing as previously described (39). Using the FLAG-PEX1^G843D^ cell line, PEX1^G843D^ tetON-PEX1^WT^ stable expression clones were generated as previously described (39). PBD fibroblasts and control lines were a gift from the Braverman Lab at McGill University. All cell lines and their derivatives were cultured in DMEM GlutaMax supplemented with 110 mg/L sodium pyruvate (Gibco #10569010), 10% fetal bovine serum (R&D Systems #FS11150H), and 100 U/mL penicillin/streptomycin (Gibco #15140122), and maintained in a humidified incubator at 37 °C and 5% CO_2_, except patient fibroblasts which were maintained identically but at 7% CO_2_ instead. For routine passaging, adherent cells were grown to ∼70 to 80% confluency, washed with Dulbecco’s phosphate buffered saline (DPBS) (Gibco #14190-144), and subsequently treated with 0.25% trypsin-EDTA (Gibco #25-200-072) for 3 to 5 min in a 37 °C incubator. Detached cells were then quenched with DMEM, trypsin removed by centrifugation, and passaged in fresh DMEM. Supernatants from cell cultures were tested routinely using the Invivogen MycoStrip mycoplasma detection kit (InvivoGen, #rep-mys) and the Lonza MycoAlert Mycoplasma detection kit (Lonza LT07-318) and *Mycoplasma spp.* contaminants were not identified.

### Hypoxia experiments

To subject cells to low oxygen conditions, cells were grown on tissue culture treated dishes for 24h prior to experimentation. The following day, culture dishes were placed in a hypoxia incubator chamber (Stem Cell #27310) with an open 10cm plate with 10mL sterile water on the lowest rack to humidify the chamber. The chamber was flushed with a mixture of 1% O_2_ and 99% N_2_ at a rate of 20 L/min for 10min, after which the sealed chamber was placed in a 37C incubator for the duration of the experiment. After treatment, the chamber was quickly disassembled, and cells lysed or fixed immediately for downstream analysis.

### Drug treatments

Cycloheximide (Sigma Aldrich #C4859) was used at a concentration of 100µg/mL. Carfilzomib (Selleck Chem #S2853) was used at 10µM. For doxycycline inducible overexpression of tetON constructs, cells were maintained in 4µg/mL of doxycycline (Sigma Aldrich #3072) for 5 days. VH298 (Sigma Aldrich #SML1896) was used at a concentration of 10µM. Roxadustat (Cayman Chemicals #15294) was used at a concentration of 50µM. Bafilomycin A1 was used at a concentration of 100nM. LYN-1604 was pre-treated for 10h at a concentration of 20µM prior to additional experimentation. CoCl_2_ was used at a concentration of 400µM.

### RNA extraction, cDNA synthesis, and qRT-PCR

Total RNA was extracted using Qiagen’s RNeasy kit (Qiagen #74106) according to the manufacturer’s instructions. Genomic DNA in the lysate was sheared using the Qiagen QIAshredder spin columns (Qiagen #79656) prior to RNA extraction. 1 μg of RNA was reverse transcribed with the iScript cDNA synthesis system (BioRad #1725035) according to the manufacturer’s protocol. The cDNA was diluted 10-fold in nuclease-free water and 0.2% was used as input for SYBR green based gene-specific quantitative PCR (BioRad #1725271). PCRs were carried out in a CFX96 Touch qPCR instrument (BioRad), and changes in gene expression were determined using the ΔΔCq method from two technical triplicates per biological replicate and a minimum of three biological replicates.

### RNA Sequencing and Gene Set Enrichment Analysis

HCT116 Pex-ZeoR cell lines harboring either NTC or PEX1 sgRNAs were detached with trypsin, pelleted, and RNA extracted using RNeasy Mini Kit (Qiagen #74104) according to manufacturer instructions, in biological triplicate. Purified RNA samples were poly-(A) enriched, reverse transcribed, and sequenced on an Illumina NovaSeq to produce paired-end 150 bp reads (Novogene). Raw fastq reads were trimmed using fastp v0.23.2 (63) alignment was performed using STAR v 2.7.11a (64), count tables were generated using htseq2 v. 2.0.2 (65) and differential expression analysis was performed using the R-package DESeq2 v. 1.40.1 (66). Differential expression comparisons were made between experimental and nontargeting CRISPRi strains in biological triplicate. Gene set enrichment analysis was performed using gseapy using genes pre-ranked by their DESeq2 Wald statistic. Enrichment was tested against the Human Hallmark 2024 gene sets from MSigDB as well as selected curated gene sets from (40–42). GSEA was performed in a single run so FDR q-values were correctly joined across the set. Gene sets with an FDR q-value ≤ 0.1 and normalized enrichment score (NES) ≥ 1.5 were considered significantly enriched. GSEA analysis scripts were written in Jupyter notebooks using Python 3.13.12. Claude Sonnet 5 was used to assist in editing version controlled and manually validated code.

### Immunoblotting

Cells were lysed directly in 2X Laemmli sample buffer, sonicated, reduced with β-mercaptoethanol, and boiled for 5 min at 95°C before loading onto BioRad TGX Stain Free 4-20% SDS-PAGE gels and transferred onto PVDF membranes using a BioRad Trans-Blot Turbo Transfer System. Membranes were blocked for 1 hr with 3% BSA in 1X TBST with 0.1% Tween20 and incubated overnight in primary antibody solutions (diluted 1:1000 in 3% BSA, 1X TBST except for anti-puromycin which was diluted 1:4000). The following day, membranes were washed 3X with 1X TBST, and incubated for 1 hr in HRP-conjugated secondary antibody solution (diluted 1:3000 anti-mouse, or 1:5000 anti-rabbit in 3% BSA, 1X TBST). Immunoreactive bands were detected using SuperSignal West Femto maximum sensitivity substrate (Thermo) and a ChemiDoc MP Imaging System (BioRad). Immunoblot signals were quantified by pixel integrated density analysis in FiJi and visualized in Prism9. Antibody for PEX1 was validated by immunoblotting of lysates from wildtype and PEX1 deficient cells. Antibody for HIF1α was validated by immunoblotting of lysates from CRISPRi HCT116dCas9 cells expressing a sgRNA for HIF1α compared to a non-targeting control sgRNA (data not shown). All other antibodies were validated using chemical perturbations with known effects. In brief, α-SCP2 was validated by immunoblotting lysates from cells with disrupted peroxisome function, α-LC3B was validated by treatment with Bafilomycin A1, α-Pro564-OH HIF1α was validated with by treatment with the hydroxylation inhibitor Roxadustat, α-phospho-eIF2α was validated by its increase in cycloheximide treatment, α-puromycin was validated by the presence of signal only in cells treated with puromycin, α-p-RS6 and α-phospho-P70S6K was validated by its decrease in rapamycin treatment (data not shown). We were unable to validate antibodies for total eIF2α, total P70S6K, and total RS6 in house since we lacked the ability to genetically deplete those factors or use small molecules for a directional effect. However, we note that these antibodies produced no secondary bands that could represent non-specific binding and are highly cited in other publications. For list of antibodies used, see Table S1.

### Immunofluorescence Staining

1.5E4 cells per well were plated on glass bottom 96-well plates (Cellvis #P96-1.5H-N) coated with 50 μg/mL poly-D-lysine (Gibco #A3890401) and fixed using 4% paraformaldehyde (Electron Microscopy Sciences #15710) in DPBS (Gibco #14190-144) for 10 minutes before washing 3X with DPBS. Cells were then permeabilized using 0.1% Triton X-100 (Thermo Fisher #A16046-AP) in DPBS for 10 minutes, washed 3X in DPBS, blocked with blocking buffer (3% BSA in 1X PBS with 0.05% Tween20) for 20 minutes, and then probed with desired antibody diluted 1:1000 in blocking buffer overnight at 4°C with gentle agitation. The following day, cells were washed 3X with DPBS and incubated with the appropriate fluorescently conjugated secondary antibody diluted 1:25000 in blocking buffer. After secondary staining, cells were counter stained for 5 min with DAPI (Invitrogen #D1306) in DPBS. Cells were then washed 3X in DPBS and stored in DPBS prior to image acquisition. For list of antibodies used, see Table S1.

### Spinning Disk Confocal Microscopy

Confocal microscopy images were acquired using an inverted spinning disc confocal microscope (Nikon Ti-Eclipse) equipped with an electron multiplying charge-coupled device camera (Fusion SN:500241) and environmental control (Okolabs stage top incubator). Image acquisition was performed with a 100X NA 1.41 oil-immersion objective. Exposure times and laser powers were standardized between replicates of an experiment. Each condition was analyzed in technical triplicate, as well as biological triplicate. Sixteen to twenty images were collected per technical triplicate (i.e., per well of a 96-well plate) in a fixed pattern. Raw micrographs were analyzed using the open-source image analysis software, CellProfiler. In brief, nuclei objects with a given radius range were identified by Otsu two class thresholding on the intensity of the DAPI channel, and the intensity of the HIF1α channel was calculated under each nucleus object. The mean integrated intensity of nuclear HIF1α was averaged across technical replicates, then across biological replicates. These data represent N>200 cells per technical replicate. Resulting analyses were plotted in Jupyter notebooks using Python 3.13.12. Claude Sonnet 5 was used to assist in editing version controlled and manually validated code.

