## Supplemental Figures for "Autophagy dependent HIF1α proteostasis is compromised in models of PEX1 deficiencies"

**Figure S1.** (A) Gene set enrichment analysis from RNA-Seq data comparing CRISPRi HCT116 cells expressing a sgRNA targeting PEX1 or a non-targeting control, with gene sets related to hypoxic or oxidative stress signaling highlighted, in addition to gene sets of established HIF1α transcriptional targets from (40–42).

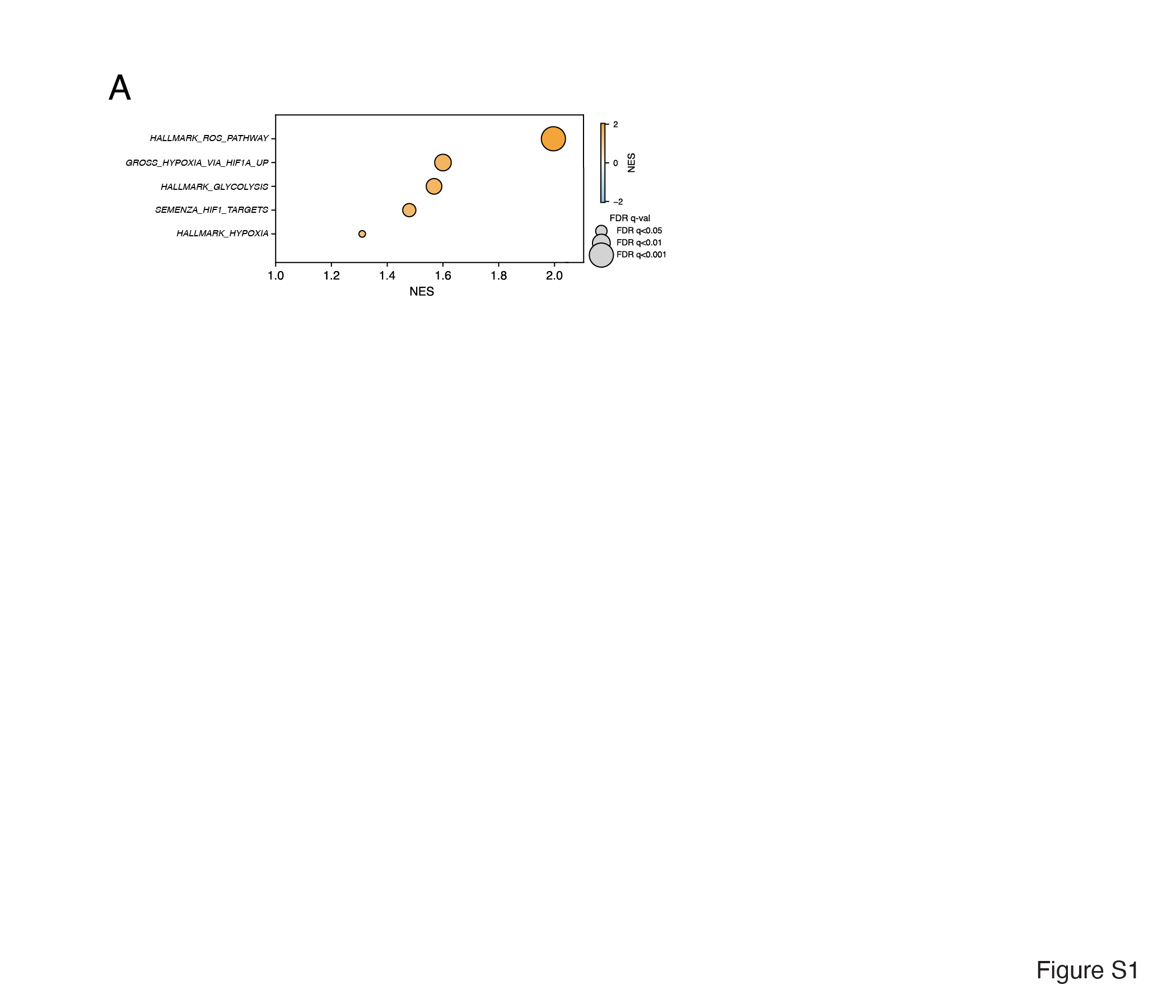

**Figure S2.** (A) Immunoblot showing levels of total and phosphorylated eIF2α as a readout of Integrated Stress Response activation in PEX1^WT^ and PEX1^G843D^ cells. (B) Pixel densitometry analysis of phosphorylated/total eIF2α from *(A)*. (C) Immunoblot of HIF1α and anti-puromycin in a puromycylation pulse labeling experiment to assess levels of global translation in PEX1^WT^ and PEX1^G843D^ cells. (D) Pixel densitometry analysis of puromycin labeling in *(C).* (E) Immunoblot showing levels of total and phosphorylated P70S6K as well as RS6 as a readout for mTORC1 signaling in PEX1^WT^ and PEX1^G843D^ cells (F) Pixel densitometry analysis of the fraction of phosphorylated P70S6K and (G) fraction of phosphorylated RS6 from *(E)* Pairwise comparisons were performed using a two-tailed unpaired Student’s t-test, where p < 0.05*, p < 0.01**, p < 0.001***

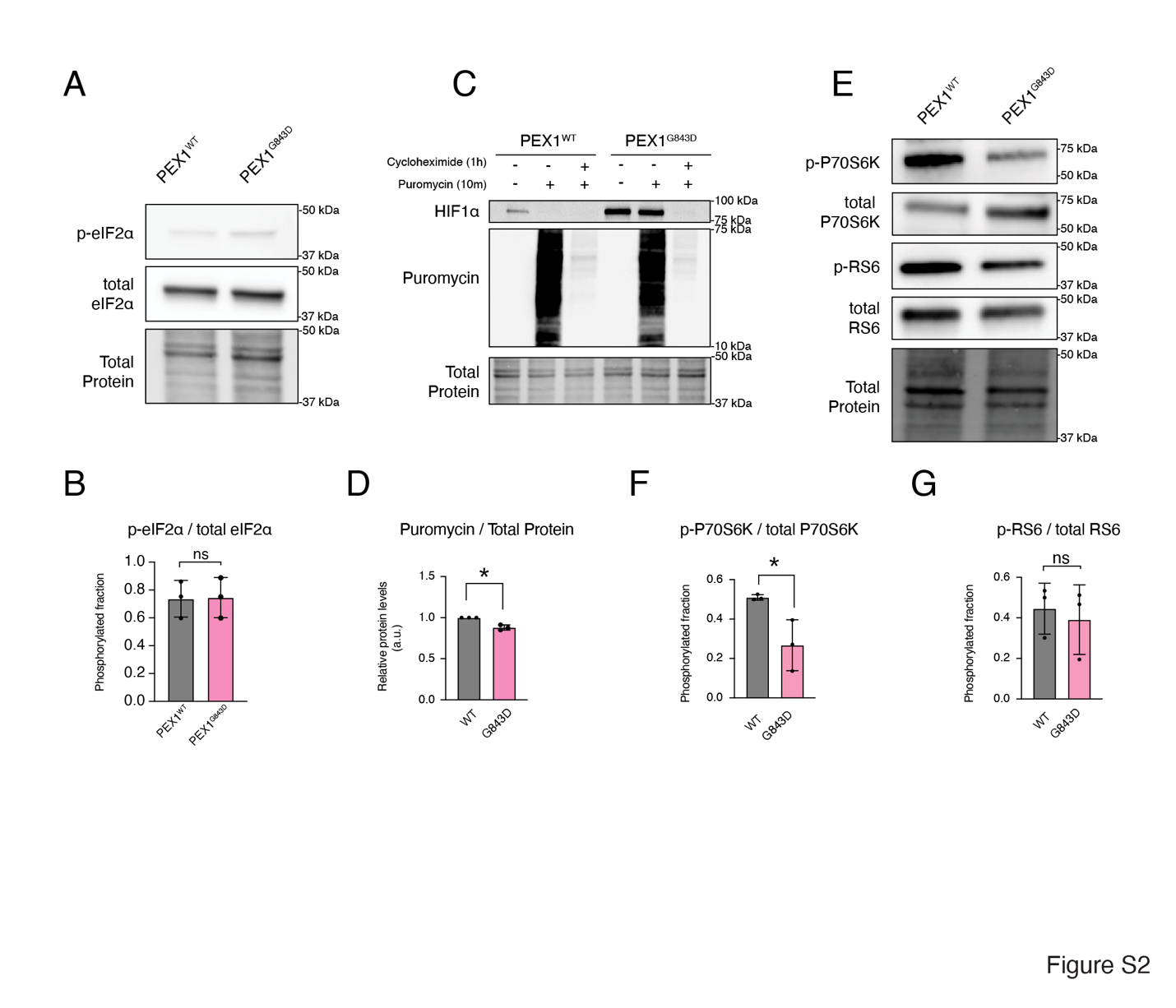

**Figure S3.** (A) Immunoblot of total HIF1α in PEX1^WT^ and PEX1^-/-^ PBD patient fibroblasts in a cycloheximide (CHX) washout experiment. Cells are first treated with CHX for 1h to allow complete degradation of HIF1α to almost zero. Then CHX is washed out and replaced with medium containing VH298 to assess the build-up of HIF1α over time when VHL dependent degradation is blocked. The SCP2 immunoblot confirms a lack of peroxisomal import in the PEX1^-/-^ fibroblasts. (B) Pixel densitometry analysis of total HIF1α levels over time from *(A).*

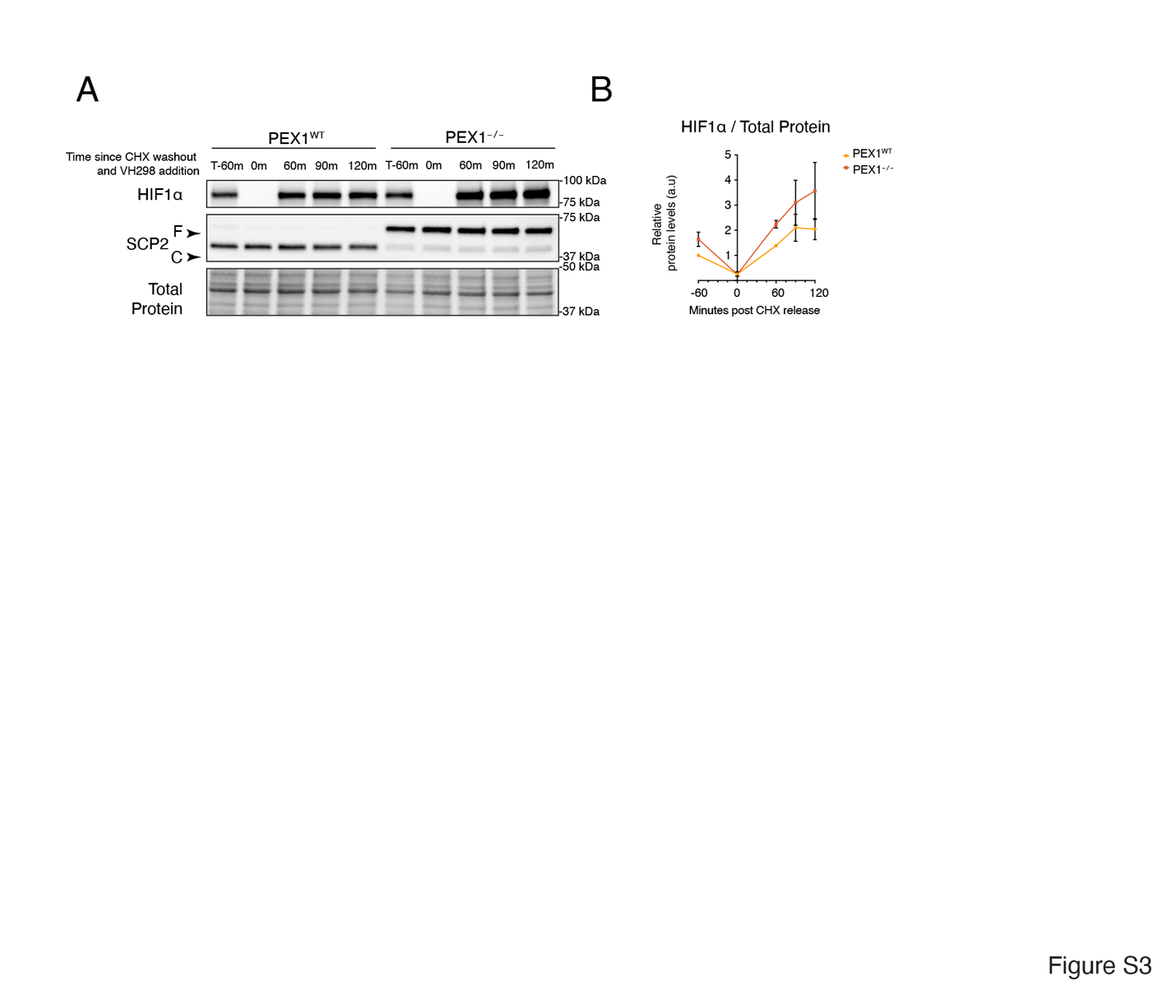
