## Supplemental Table 1 for "Autophagy dependent HIF1α proteostasis is compromised in models of PEX1 deficiencies"

**Table S1. List of Antibodies used for western blot and immunofluorescence**

| Antibody Name | Catalog # | Supplier | Dilution Used | Use |
| --- | --- | --- | --- | --- |
| PEX1 | 611719 | BD Biosciences | 1:1000 | Western Blot |
| SCP2 | HPA027317 | Sigma/Atlas | 1:1000 | Western Blot |
| HIF1α | 610958 | BD Biosciences | 1:1000 | Western Blot |
| HIF1α | 36169 | Cell Signaling | 1:1000 | Immunofluorescence |
| Pro564-OH-HIF1α | 3434 | Cell Signaling | 1:1000 | Western Blot |
| HIF2α | 59973 | Cell Signaling | 1:1000 | Western Blot |
| Phospho-eIF2α (S51) | 9721 | Cell Signaling | 1:1000 | Western Blot |
| Total-eIF2α | 9722 | Cell Signaling | 1:1000 | Western Blot |
| Puromycin | MABE342 | Sigma-Aldrich | 1:5000 | Western Blot |
| Phospho-P70S6K (T389) | 9234 | Cell Signaling | 1:1000 | Western Blot |
| Total-P70S6K | 9202 | Cell Signaling | 1:1000 | Western Blot |
| Phospho-RS6 (S235/236) | 4857 | Cell Signaling | 1:1000 | Western Blot |
| Total-RS6 | 2217 | Cell Signaling | 1:5000 | Western Blot |
| LC3B | 2775 | Cell Signaling | 1:2000 | Western Blot |
| AF647 goat anti-rabbit IgG [H+L] | A-21245 | Invitrogen | 1:2500 | Immunofluorescence |
| AF594 goat anti-mouse IgG [H+L] | A-11032 | Invitrogen | 1:2500 | Immunofluorescence |
| Goat anti-rabbit HRP | 1706515 | BioRad | 1:5000 | Western Blot |
| Goat anti-mouse HRP | 1706516 | BioRad | 1:3000 | Western Blot |
